# K-mer counting with low memory consumption enables fast clustering of single-cell sequencing data without read alignment

**DOI:** 10.1101/723833

**Authors:** Christina Huan Shi, Kevin Y. Yip

## Abstract

K-mer counting has many applications in sequencing data processing and analysis. However, sequencing errors can produce many false k-mers that substantially increase the memory requirement during counting. We propose a fast k-mer counting method, CQF-deNoise, which has a novel component for dynamically identifying and removing false k-mers while preserving counting accuracy. Compared with four state-of-the-art k-mer counting methods, CQF-deNoise consumed 49-76% less memory than the second best method, but still ran competitively fast. The k-mer counts from CQF-deNoise produced cell clusters from single-cell RNA-seq data highly consistent with CellRanger but required only 5% of the running time at the same memory consumption, suggesting that CQF-deNoise can be used for a preview of cell clusters for an early detection of potential data problems, before running a much more time-consuming full analysis pipeline.

## Introduction

The high-throughput nature and ever decreasing cost of next-generation sequencing (NGS) technologies have enabled the development of experimental methods for a variety of applications [1, 2]. For applications that produce a large number of sequencing reads, such as whole-genome sequencing for de novo assembly [3], directly operating on the sequencing reads could be slow and memory-prohibitive when the depth-of-coverage is high. As a result, it has become common to summarize the sequencing data by the list of all k-mers (i.e., length-k sub-sequences) and their occurrence frequencies in the sequencing reads. These k-mer counts are useful for various downstream tasks, including error correction [4–6], de Bruijn graph construction [6–8], read clustering and query [9], and genome size estimation [10, 11].

In k-mer counting, the major computational concerns include counting and querying efficiency, and memory consumption. These issues are particularly critical when the sequenced genome is large, the depth-of-coverage is high, or when there is a limited amount of memory available. Many k-mer counting methods have been proposed to deal with these concerns.

In terms of counting efficiency, one common way to speed up the counting process is multi-threading. The main bottleneck of multi-threading methods is the overhead caused by locking, and different methods have used different approaches to tackle it. For example, Jellyfish2 [12] takes advantage of the CAS (compare and set) assembly instruction to build a lock-free hash table for parallelism. As another approach, Squeakr [13] allows each thread to store values of shared variables in a local data structure temporarily, which will later be merged into the global data structure.

In terms of memory consumption, one way to handle large data sets is to process only a subset of data in memory at a time. For example, DSK [14] and KMC [15] use the disk to split a large data set into different bins (files), and process each bin separately one by one. When the intermediate results of the different bins are ready, they are further combined to produce the final k-mer counts.

Another way to reduce memory consumption while allowing efficient k-mer queries is to use AMQ (Approximate Membership Query) data structures. These structures are for querying whether a particular object (a k-mer in this case) is contained in a set/multi-set or not. If the set contains the object, the query result is always positive, and thus these structures guarantee no false negatives. On the other hand, if the set does not contain the object, there is a certain probability that the query result would still be positive, and the rate of such false positives is determined by properties of the AMQ data structure.

One widely used AMQ data structure is the Bloom filter, which is a bit vector for recording the objects stored. Each object to be stored is fed to multiple independent hash functions to generate as its signature a list of hash values, with each value occupying one bit of the Bloom filter with equal probability. The Bloom filter for a set of objects is simply the disjunction of the hash values of all its objects. To query whether an object is contained in the set, its hash value signature is computed. If any of the positions occupied by the hash values is not set in the Bloom filter, the object is definitely not contained in the set. Otherwise, the object is either contained in the set or it is not contained in the set but its hash values form a subset of the hash values of the objects in the set. The probability of the latter, false positive case (also called a “false drop” in the literature) is controlled by the size of the Bloom filter, the number of hash functions used, and the number of objects in the set. Assuming that each bit in the Bloom filter is set independently, the false positive probability is 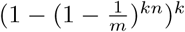 [16], which is approximately (1 − *e*^*−kn/m*^)^*k*^ when *m* is large, where *m* is the number of bits in the Bloom filter, *k* is the number of hash functions, and *n* is the total number of unique objects inserted into the Bloom filter.

Some studies have attempted to improve the speed efficiency of the Bloom filter. For example, blocked Bloom filter [17] divides up a Bloom filter into multiple smaller Bloom filters each of which can fit into the cache line, which improves the cache efficiency at the expense of a higher false positive rate.

To use the Bloom filter for object counting, one way is to use *c* bits for each data slot (instead of one bit) to record how many times this slot is set, which forms a counter with a value up to 2^*c*^ − 1 [18]. This variation of the Bloom filter permits object removal, by decrementing the counters in the *k* corresponding slots of the object to be removed, again with a chance of false decrements due to the false positive rate. A drawback of this counting approach is that every data slot is allocated *c* bits regardless of the number of times that it is set, leading to unnecessary memory consumption. Besides, it does not provide an easy way to query the occurrence count of an object because each counter is associated with a slot and each slot can be shared by many objects. Some other variations of the counting Bloom filter have been proposed in the literature [19], including ones that allow the query of object occurrence counts [20].

The counting quotient filter (CQF) [19] is a recently proposed AMQ data structure for object counting that is more space efficient than the counting Bloom filter, and it allows dynamic re-sizing of the data structure as more data are added. We explain CQF in detail in Materials and Methods.

The ideas described above can also be combined. For example, Squeakr [13] implements a multi-threaded CQF, which can perform k-mer counting with both time and space efficiency.

A practical issue of k-mer counting is the presence of errors in the sequencing reads, which increases the number of unique k-mers by creating false k-mers that do not actually exist in the original sequences. Since the number of distinct random errors grows with the number of sequencing reads produced, the memory consumption of k-mer counts can keep increasing with sequencing depth even though the number of true k-mers in the original sequences stays constant.

An important property of false k-mers is that they usually have much lower occurrence counts than the true k-mers due to the random nature of most sequencing errors. Therefore, some previous k-mer counting methods simply assume low-frequency k-mers are errors and discard them. For example, BFCounter [21] and Turtle [22] use a Bloom filter to detect whether a k-mer has been encountered before and a separate data structure for the actual counting of k-mers that appear at least twice in the sequencing reads. In this way, the singletons (k-mers that appear only once in the sequencing reads) will not take up space in the second data structure. Alternatively, Jellyfish2 [12] filters low-frequency k-mers by keeping a counting Bloom filter with a small number of bits per slot, and when the counter reaches a pre-defined threshold (such as two), further k-mer counting will be performed using another data structure that is more space efficient.

We argue that these approaches to handling false k-mers are not ideal in two aspects. First, false k-mers take up space in the data structures during the counting process, with a possibility of using up all the available memory before the whole counting process is complete. Therefore, they should be removed as early as possible. Second, using an extra step to identify singletons leads to extra overheads in terms of running time and possibly memory consumption. It would be more preferable to combine counting and false k-mer removal in the same process.

In this paper, we propose a method called CQF-deNoise that uses the CQF data structure for counting k-mers while removing false k-mers on the fly. Based on a user-specified wrong removal tolerance threshold, CQF-deNoise automatically determines the suitable time and number of rounds of false k-mer removal. As a result, the number of unique k-mers in the CQF during the whole counting process remains largely constant and is much smaller than the total number of unique true and false k-mers. We show that as compared to several state-of-the-art k-mer counting methods, CQF-deNoise consumes less memory, runs competitively fast, but at the same time gives k-mer counts that are highly accurate. We also show that the fast and low-memory k-mer counting achieved by CQF-deNoise makes it possible to cluster cells based on single-cell RNA-seq data using only 5% of the time required by a standard pipeline while the clusters produced remain highly similar.

## Results

The details of CQF-deNoise are given in Materials and Methods. We tested its performance and compared it with several other k-mer counting methods using four data sets with diverse properties (Table 1). All methods were tested on the same machine, and parameter values of CQF-deNoise were determined by an automatic procedure described in Materials and Methods with the actual values used listed in Table 2.

**Table 1.** Data sets used for testing the performance of CQF-deNoise and comparing it with other k-mer counting methods. Data sets were chosen from four species with very different genome sizes. The sequencing data also had different depths of coverage of the genomes.

| Species | Reference genome | Genome size (Mb) | Data sets (SRA accessions) | Total read length (Gb) | Depth-of-coverage |
| --- | --- | --- | --- | --- | --- |
| C. elegans | WBcel235 | 100 | SRR7693585-7693591 | 29.0 | 290× |
| F. vesca | FraVesHawaii.1.0 | 240 | SRR072005-072014,<br>072029, 5275218, 5799056-5799057 | 14.1 | 59× |
| Z. mays | B73 RefGen_v4 | 2,100 | SRR7753852-7753853,<br>7753855, 7753858-7753861,<br>7753878-7753884 | 128.8 | 61× |
| H. sapiens | GRCh38.p12 | 3,257 | SRR2831527-2831541 | 199.6 | 61× |

**Table 2.** Parameter values of CQF-deNoise. The length of hash value signature (*p*), quotient (*q*) and remainder (*r*) automatically determined by our algorithm based on the number of rounds of k-mer removal (*m*) used in our empirical tests.

| $m$ | $p$ | $q$ | $r$ |
| --- | --- | --- | --- |
| 0,1 | 40 | 32 | 8 |
| 2 | 39 | 31 | 8 |
| 5,10,15 | 38 | 30 | 8 |
| 20,25,30,40,50 | 37 | 29 | 8 |

### CQF-deNoise has a low wrong removal rate of true k-mers

Since CQF-deNoise removes potential false k-mers based on their low occurrence frequencies, it could accidentally remove some true k-mers. We tested how many true k-mers (those that appear in the reference genome) were wrongly removed by CQF-deNoise using the C. elegans data set, which had the highest depth of coverage and thus the highest expected false-to-true k-mer ratio in the data caused by the sequencing errors.

When we did not perform noise removal, the frequency distribution of the full set of k-mers was clearly multi-modal (Figure 1a), with a peak at around 160 that likely corresponded to the median coverage of the true k-mers, and another peak at 1 that should contain mostly false k-mers. Since a local minimum was observed at 50, we used it as the demarcation point and manually set the number of noise removal rounds of CQF-deNoise for each value from 0 to 50 to inspect the change of counting results.

**Figure 1.**
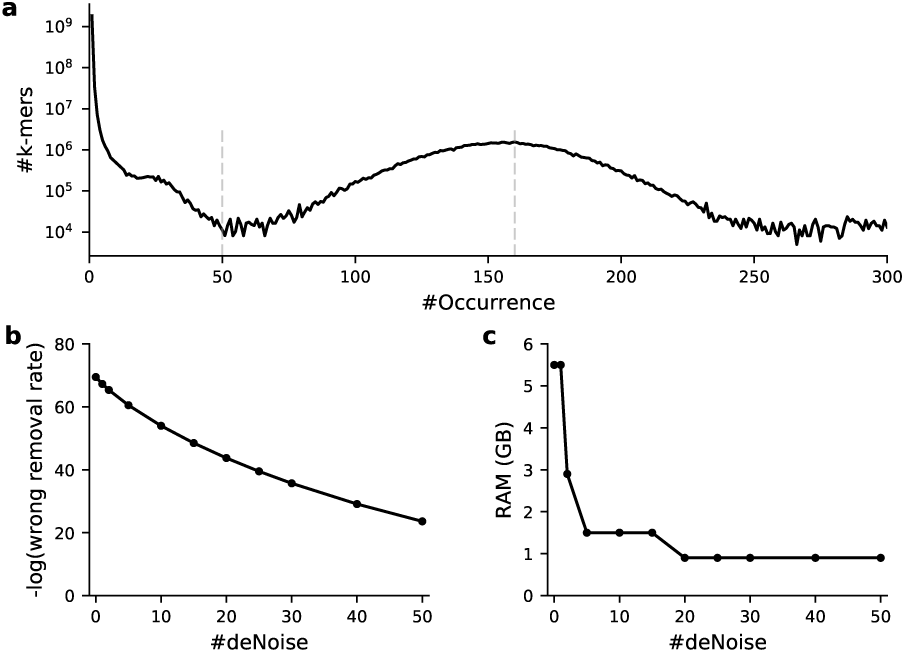
Trade-off between memory consumption and wrong removal of true k-mers based on the C. elegans data set with *k* = 28. (a) Distribution of k-mer occurrence counts without noise removal. (b) Estimated wrong removal rate and (c) memory consumption at different numbers of k-mer removal rounds. The estimated wrong removal rate was defined as the fraction of true k-mers having an occurrence count no larger than the number of rounds of noise removal, estimated by the Poisson distribution.

As the number of noise removal rounds increased, as expected more and more true k-mers were wrongly removed, but the rate remained low, reaching the maximum of only 10^*−*23.6^ with 50 rounds of noise removal (Figure 1b), which is negligible given the high sequencing depth (290×). As to be discussed below, in real situations the number of noise removal rounds determined automatically by CQF-deNoise is usually much smaller than 50, and thus the fraction of true k-mers being removed is very small in practice.

At the same time, the memory consumption rapidly dropped from 5.5GB to 0.9GB as the number of noise removal rounds increased from 0 to 20, which was then stabilized thereafter (Figure 1c). To interpret this memory consumption objectively, we performed the following conceptual analysis. Assuming that most true k-mers in the genome are unique and there is a uniform read coverage of the whole genome in the data set, there would be around 100 million unique true k-mers in the C. elegans genome, each with an occurrence count of 160. Since two counting slots are required to store this average occurrence count when the remaining parameter, *r*, is equal to 8 in addition to a slot storing the remainder (Materials and Methods; Table 2), the total number of slots in the CQF would be at least 300 millions. Further, since the number of slots in the CQF has to be a power of 2, the smallest number of slots becomes 2^29^, which was exactly the number of slots in the CQF we constructed, showing that our memory usage of 0.9GB was optimal in this case.

### CQF-deNoise has high counting accuracy

We evaluated the counting accurcy of CQF-deNoise using the C. elegans data set in two ways. First, since singleton k-mers are mostly false k-mers, we evaluated the proportion of singleton k-mers that remained in the CQF (the “singleton survival rate”) and the proportion of non-singleton k-mers that did not remain in the CQF (the “non-singleton removal rate”). Singletons can survive due to the intrinsic nature of CQF, that a low-occurrence k-mer can happen to share the same hash value as one or more other k-mers, and thus their counts add up. A non-singleton can be removed if its occurrence count is no more than one every time when a round of noise removal starts. From the results, the singleton survival rate remained fairly stable across the different numbers of noise removal rounds, with only around 0.1%-0.2% of the singletons remained in the CQF at the end of counting (Figure 2a). For the non-singletons, less than 2.5% of them were removed (Figure 2b), and many of these non-singletons are expected to be false k-mers based on the shape of the bimodal k-mer count distribution (Figure 1a).

**Figure 2.**
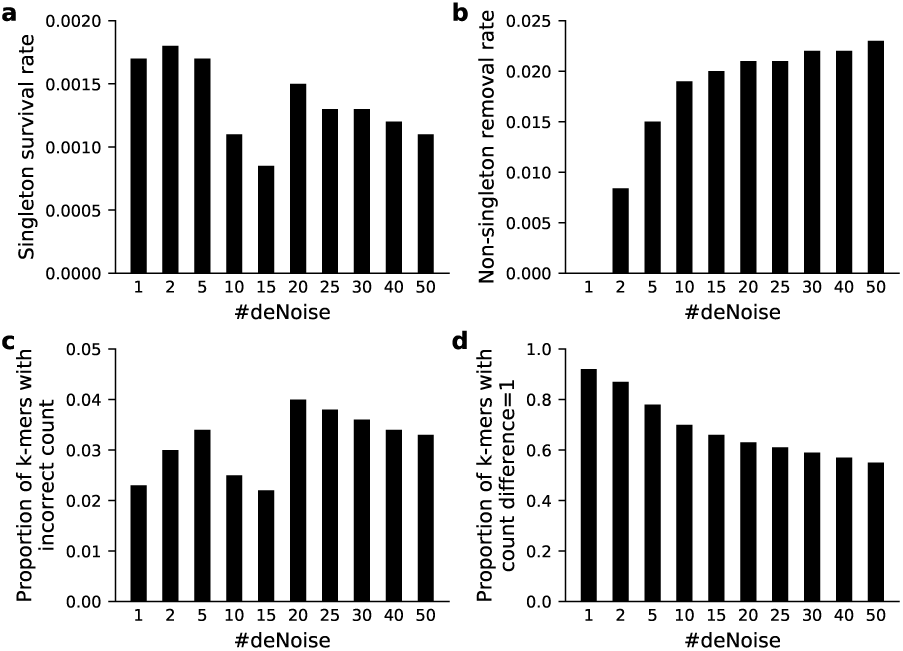
Counting accuracy based on the C. elegans data set. (a) Singleton survival rate, (b) non-singleton removal rate, (c) proportion of k-mers in CQF with a count different from its real count, and (d) among all the k-mers in CQF with count different from its real count, the proportion of k-mers with count difference being 1, with different numbers of rounds of noise removal.

Second, for the k-mers that remained in the CQF at the end of counting, we evaluated the correctness of their counts. We observed that only 2%-4% of these k-mers had an incorrect count (Figure 2c), and among these k-mers with an incorrect count, more than half of them had a difference of only 1 between the actual count and the count in the CQF (Figure 2d), showing that the noise removal procedure of CQF-deNoise had minimal effects on counting accuracy.

### CQF-deNoise uses less memory than other k-mer counting methods

To benchmark the computational performance of CQF-deNoise, we compared it with four state-of-the-art k-mer counting methods, namely BFCounter [21], Jellyfish2 [12], KMC3 [15], and Squeakr [13]. BF-Counter, Jellyfish2 and Squeakr are memory-based, while KMC3 was designed to be a disk-based method, although it also provided an in-memory mode, which we used in our comparisons.

We first compared the memory consumption of the different methods based on the four data sets. From the results (Figure 3a,b), CQF-deNoise consistently consumed the smallest amount of memory on all four data sets for both values of *k* tested. Compared to the next method with the least memory consumption, CQF-deNoise had a memory usage reduction from 49% (F. vesca data set, *k* = 55) to 76% (C. elegans data set, *k* = 28).

**Figure 3.**
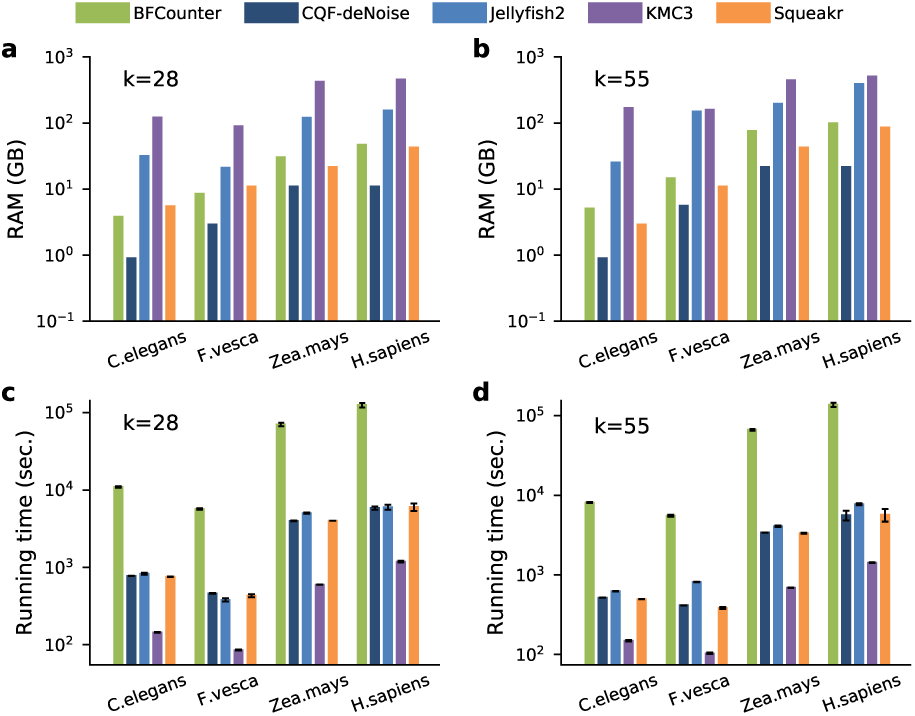
Comparison of the k-mer counting methods. Memory usage when (a) k=28 and (b) k=55. Running time when (c) k=28 and (d) k=55.

The running time of CQF-deNoise was similar to that of Jellyfish2 and Squeakr and was consistently faster than BFCounter (Figure 3c,d). It was not faster than KMC3, but the short counting time of KMC3 came with the cost of a much higher memory consumption, which was 21 (Z. mays data set, *k* = 55) to 187 (C. elegans data set, *k* = 55) times of the memory consumption of CQF-deNoise.

To evaluate the scalability of the different methods, we considered the C. elegans data set (which had the highest depth-of-coverage), and sub-sampled reads at different depths. The results show that CQF-deNoise used the least amount of memory at all depth values (Figure 4a). Importantly, the memory consumption increase with respect to sequencing depth was smallest for CQF-deNoise, since it was largely unaffected by the increasing amount of false k-mers in the data. In terms of running time, the order of the different methods remained unchanged at the different depths of coverage (Figure 4b).

**Figure 4.**
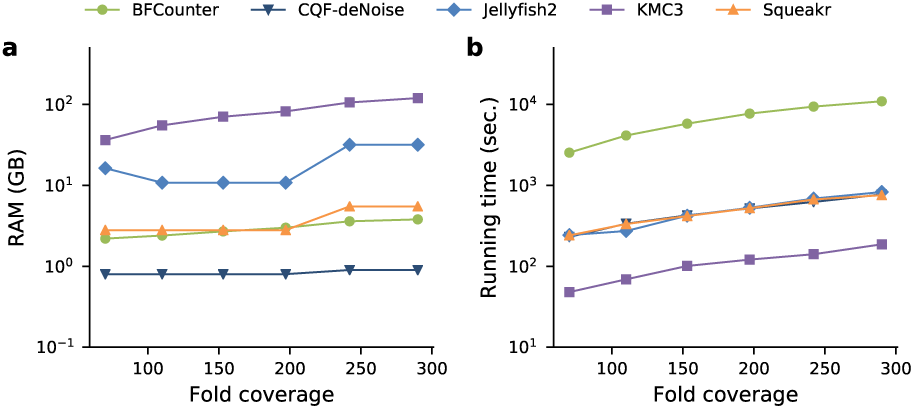
Scalability of the k-mer counting methods (a) Memory consumption and (b) running time of the different methods on random subsets of reads from the C. elegans data set at 70*×* to 290*×* coverage.

These results show that CQF-deNoise performs k-mer counting with lower memory consumption while running as fast as the other memory-based methods.

### K-mer count querying using CQF-deNoise is efficient

Another important performance indicator is the time needed to query the occurrence counts of k-mers after the counting process, when all the counts are already loaded into memory. We compared the different methods using the C. elegans and F. vesca data sets only due to the large amount of time needed by some of the published methods to query from the other two data sets. For each of the two data sets, we queried both k-mers that existed in the sequencing reads and were contained in all the counting data structures as well as k-mers that did not exist in the reads.

The results (Table 3) show that CQF-deNoise completed these queries using the smallest amount of time among all the methods for both data sets and for both types of k-mers. The high query efficiency of CQF-deNoise was likely due to the compactness of its data structure.

**Table 3.**
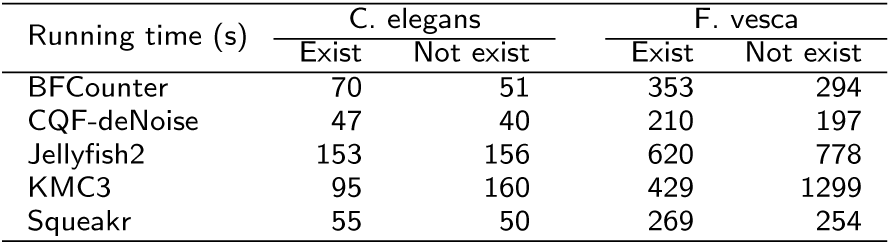
Query performance of the different methods. For each species, the running time of querying a) random k-mers that existed in the sequencing reads and were contained in the data structures (“Exist”) and b) random k-mers that did not exist in the sequencing reads (“Not exist”) are reported. Each query set contained approximately 100 million and 550 million k-mers in the case of C. elegans and F. vesca, respectively.

| Running time (s) | C. elegans |  | F. vesca |  |
| --- | --- | --- | --- | --- |
|  | Exist | Not exist | Exist | Not exist |
| BFCOUNTER | 70 | 51 | 353 | 294 |
| CQF-deNoise | 47 | 40 | 210 | 197 |
| Jellyfish2 | 153 | 156 | 620 | 778 |
| KMC3 | 95 | 160 | 429 | 1299 |
| Squeakr | 55 | 50 | 269 | 254 |

### Cell clusters can be accurately identified by k-mer counts in single-cell RNA-seq data

Finally, we explored the clustering of single cells as an application of k-mer counts. We obtained a single-cell RNA-seq (scRNA-seq) data set of peripheral blood mononuclear cells produced using the 10x Chromium platform [23]. For each cell, we computed the occurrence counts of either all k-mers or only the k-mers that appear in annotated protein-coding transcripts using CQF-deNoise. In both cases, two-dimensional projection of the cells based on these k-mer counts using UMAP [24] showed that there were four cell clusters (Figure 5a,b). Using SingleR [25] to annotate the cell types, we found that monocytes and B-cells formed two distinct clusters, while the other two clusters contained a mixture of cells from different types with the natural killer (NK) cells occupying a corner of one of these clusters.

**Figure 5.**
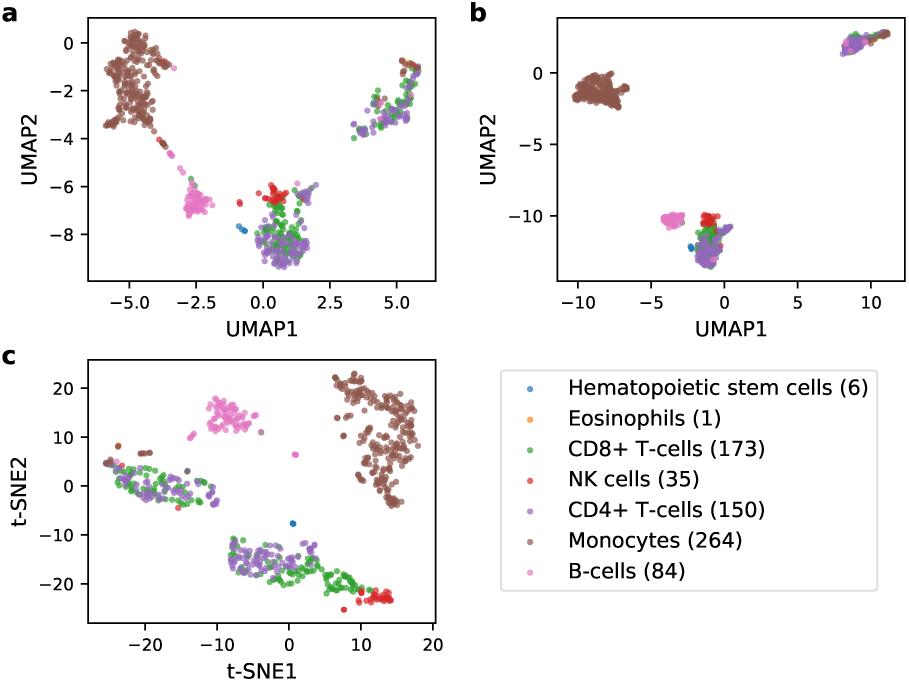
Visualization of single cells based on the scRNA-seq data set. (a,b) Two-dimensional UMAP projection of single cells based on the counts of all k-mers (a) or only the k-mers in annotated protein-coding transcripts (b). (c) Two-dimensional projection of single cells produced by CellRanger. In the legend, the numbers in brackets represent cell numbers of the different types.

To evaluate the quality of these clusters, we applied CellRanger [26], a pipeline designed for 10x scRNA-seq data, to this data set. The two-dimensional projection also showed four clusters with very similar cell type distributions as the ones produced by CQF-deNoise (Figure 5c).

To have a more quantitative comparison between CQF-deNoise and CellRanger, we used k-means [27] to produce four clusters using either the protein-coding transcript k-mer counts or the transcript expression levels computed by CellRanger. The adjusted Rand index (ARI) between the two sets of clusters was high (0.94), while when they were individually compared with the SingleR annotations, the ARI values were very close (both 0.60), showing that the two sets of results were highly consistent.

The advantage of using k-mer counts over a full analysis pipeline that includes read alignments lies in the computational resources required. When we did not restrict the memory consumption, the CQF-deNoise pipeline was completed in 19 minutes using 6.4GB of memory, while CellRanger used 72 minutes (3.8 times of CQF-deNoise) and 142.7GB of memory (22.3 times of CQF-deNoise). When we set the maximum memory usage to 7GB, CellRanger took 374 minutes to complete, which was 19.7 times of CQF-deNoise; or equivalently, CQF-deNoise only used 5% of CellRanger’s running time.

These preliminary results suggest that k-mer counts can potentially be used to provide a quick preview of the analysis results that a full analysis pipeline would take a much longer time to produce.

## Discussion and conclusion

In this paper, we have proposed a memory-efficient k-mer counting method, CQF-deNoise, that uses less memory than other stat-of-the-art k-mer counting methods but runs as fast as the memory-based methods. Its low memory consumption is attributed to a procedure that removes potential false k-mers caused by sequencing errors. When compared to the next method with the lowest memory consumption, CQF-deNoise used 49%-76% less memory.

The false k-mer removal procedure of CQF-deNoise automatically determines the time and number of rounds of k-mer removal based on a user-specified maximum tolerable rate of wrongly removing true k-mers. We have shown that this procedure was effective in removing false k-mers while the counts of true k-mers were only minimally affected. Although some other k-mer counting methods can also remove some potential false k-mers, they do it in a fairly arbitrary way by for example removing all singleton k-mers at the end of counting, resulting in an increase of memory usage as the sequencing depth (and thus number of false k-mers) increases. In comparison, the dynamic removal procedure of CQF-deNoise determines the k-mers to remove based on properties of the sequencing data, and it led to virtually constant memory usage within a range of sequencing depths.

We demonstrated that k-mer counts can be used to cluster single cells based on their scRNA-seq profiles, with the clusters formed highly consistent with the ones formed by a full analysis pipeline for scRNA-seq data but our k-mer-based approach used much less running time and/or memory. Conceptually, as long as a task requires only a distance matrix of single cells as input, such as various types of visualization, clustering and trajectory inference, our k-mer-based approach can be used. Whether there are some particular data types that require filtering or normalization steps that cannot be performed efficiently using CQF-deNoise needs to be investigated in more detail.

We provide CQF-deNoise as an open-source package. The main program for counting k-mers contains additional options for users who want to have more control of the counting process, such as specifying the exact number of noise removal rounds to be performed. The package also comes with a number of extra tools for manipulating CQFs in general.

One limitation of CQF-deNoise is that some k-mers that remain in the CQF can actually have a lower occurrence count than some k-mers completely removed from the CQF. This is because whether a k-mer would be removed depends not only on its occurrence count, but also on the moments of its different occurrences. A k-mer remains if and only if there are two or more occurrences of it within any of the time periods between two noise removal rounds. This would not affect the removal of singleton k-mers, but by chance some false k-mers do occur more than once and are not removed in this way. When it is crucial to have as few false k-mers remaining in the CQF as possible, one possible remedy is to construct a histogram of k-mer occurrence frequencies after counting, use it to determine occurrence counts that likely belong to the false k-mers by inspecting the distribution (as was done in Figure 1), and finally perform a post-processing round of noise removal using the identified threshold.

## Materials and Methods

### The counting quotient filter

CQF-deNoise uses CQF [19] for efficiently computing the occurrence frequencies of k-mers, with a novel denoise method for removing k-mers that likely occur due to sequencing errors. Here we first explain how CQF works and how we implemented it, and then we will describe our de-noise method in the next subsection.

CQF is an AMQ data structure for object counting. It represents a multiset *S* by using a hash function *h* to map every object *x* to a *p*-bit representation, where *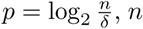*, *n* is the expected maximum number of distinct objects in *S*, and *δ* is the desired false positive query rate. Unlike the Bloom filter that directly uses all *p* bits as the signature of an object, in CQF, the first *q* bits (called the quotient) are used to determine the canonical memory slot (called the “home slot”) that an object should be stored in, and the remaining *r* = *p−q* bits (called the remainder) are actually stored in the memory slot to indicate that the slot contains an object with that signature. The CQF thus contains a table of 2^*q*^ data slots each storing *r* bits of data. When collision occurs, i.e., when an object’s home slot has already been taken by another object, the exact memory slot to be used for storing it is determined by a variant of linear probing, with the objects assigned to adjacent slots following the same order as their hash values. Object insertion, query and deletion are all assisted by some additional metadata that contain information about whether the CQF has stored any object with a particular quotient and where each run of data slots of objects with the same quotient ends.

Object counts are stored in CQF in three different ways according to their values. For an object that has appeared only once, no additional information is stored, which serves as an indicator that the object count is one. For an object that has appeared twice, an additional slot is assigned right after the original slot, and it also stores the same remainder value of the object. For an object that has appeared three or more times, right after the original slot assigned to the object, one or more additional slots are assigned as the counter of the object, followed by another slot storing the remainder of the object again to signify the end of the counter. The number of slots assigned to the counter can be dynamically modified, such that CQF can handle object counts of very different magnitudes at the same time. To distinguish between a slot storing a remainder and one storing a counter, the first slot assigned as part of the counter must have a value smaller than the remainder of the object. This is sufficient for indicating that the slot is a counter, because by definition objects in the same run are stored in ascending order of their remainder values. The encoding scheme for counters also employs some additional rules to make sure that all combinations of remainder and counter values can be stored and retrieved correctly (to be explained below).

### Our implementation of CQF, with a more space-efficient counter encoding scheme

We adopted the implementation of CQF in Squeakr [13] on the basis of its efficient C++ code and multi-threading option, but we made two custom designs. First, since our CQF was used for counting DNA sequences, we chose the ntHash function [28], which was specifically designed for nucleotide sequences and was shown to perform better than several mainstream hash functions. Second, we proposed an alternative encoding scheme for the counters, which combines ideas from both the original encoding scheme used in CQF and the Most-Significant Bit (MSB) encoding scheme. The basic idea is to use the most significant bit to indicate whether this counter still occupies additional slot(s), rather than storing the remainder of the object again after the last counter slot.

Specifically, suppose the occurrence count of an object with remainder *x* is *C*, and each remainder occupies *r* bits. As in the case of the original encoding scheme of CQF, how the occurrence count is stored in CQF-deNoise depends on the values of *C* and *r*.

If *C* = 1, a single slot is assigned that stores *x* as its value. If *C >* 1, multiple slots are assigned, with the first slot storing *x* as its value and the remaining slots storing an encoding of the counter. Since such counter slots are used only when *C >* 1, we store *C −* 1 instead of *C*. To determine how *C* − 1 is represented, we convert it into binary form and count the number of bits required. Suppose *b* bits are needed, then *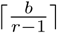* slots are allocated, and the last *r −* 1 bits of each of these slots together store the binary form of *C −* 1. After that, for all the counter slots except the last one, the most significant bit among its *r* bits is set to 1, to indicate that it is not the last slot of this counter. Finally, if the number stored in the first of these counter slots is larger than *x*, a slot storing the value zero is added at the beginning as an additional escape value.

There are several major differences between the original encoding scheme of CQF and the encoding scheme of CQF-deNoise:

- CQF stores *x* both before and after the counter slots, while CQF-deNoise only stores it before the counter slots. Instead, every counter slot reserves the most significant bit to indicate whether there are more counter slots to come or not.
- CQF reserves the values 0 and *x* for special meanings, such that counter values need to be encoded in a way that depends on *x*. In contrast, counter values in CQF-deNoise can be easily determined by simply ignoring the most significant bit of each counter slot.
- CQF needs to specially handle the case *x* = 0 since it requires putting a value smaller than *x* in the first counter slot. CQF-deNoise does not need to consider *x* = 0 as a special case, since it only requires putting a value not larger than *x* (rather than smaller than *x*) in the first counter slot.

Table 4 shows some examples of how CQF and CQF-deNoise encode the counter values.

**Table 4.**
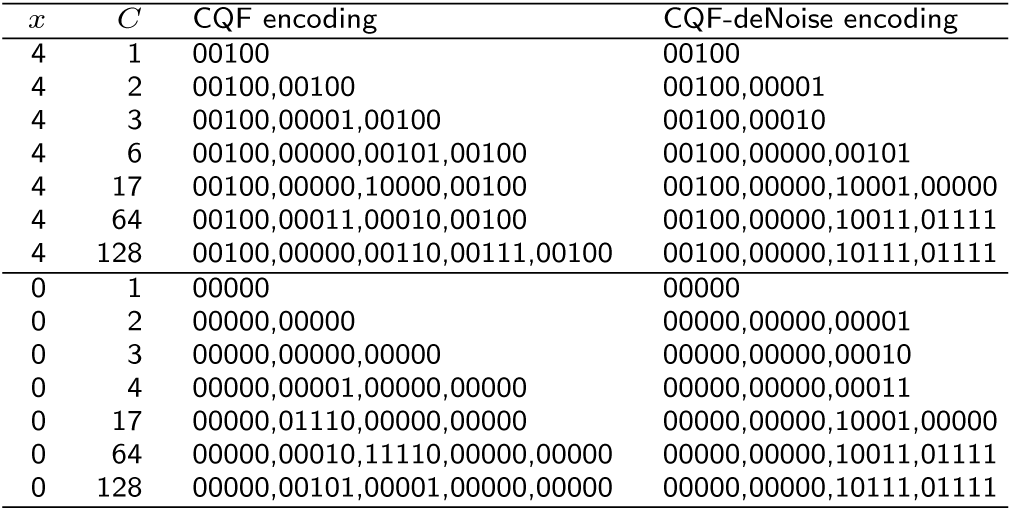
Encoding examples of CQF and CQF-deNoise. This table contains examples that show how CQF and CQF-deNoise encode the occurrence count *C* of an object with remainder *x* containing *r* = 5 bits.

| $x$ | $C$ | CQF encoding | CQF-deNoise encoding |
| --- | --- | --- | --- |
| 4 | 1 | 00100 | 00100 |
| 4 | 2 | 00100,00100 | 00100,00001 |
| 4 | 3 | 00100,00001,00100 | 00100,00010 |
| 4 | 6 | 00100,00000,00101,00100 | 00100,00000,00101 |
| 4 | 17 | 00100,00000,10000,00100 | 00100,00000,10001,00000 |
| 4 | 64 | 00100,00011,00010,00100 | 00100,00000,10011,01111 |
| 4 | 128 | 00100,00000,00110,00111,00100 | 00100,00000,10111,01111 |
| 0 | 1 | 00000 | 00000 |
| 0 | 2 | 00000,00000 | 00000,00000,00001 |
| 0 | 3 | 00000,00000,00000 | 00000,00000,00010 |
| 0 | 4 | 00000,00001,00000,00000 | 00000,00000,00011 |
| 0 | 17 | 00000,01110,00000,00000 | 00000,00000,10001,00000 |
| 0 | 64 | 00000,00010,11110,00000,00000 | 00000,00000,10011,01111 |
| 0 | 128 | 00000,00101,00001,00000,00000 | 00000,00000,10111,01111 |

From Table 4, we can see that the encoding scheme of CQF-deNoise rarely uses more slots than the scheme of CQF. In fact, by not having the remainder stored twice before and after the counting slots, the encoding scheme of CQF-deNoise usually consumes less space. Table 5 compares the number of slots required by the two encoding schemes for all possible occurrence counts within two practical ranges in genomic applications. In these two settings, the encoding scheme of CQF-deNoise consumes less memory in 61% and 22% of the cases, respectively, while it consumes more memory in only 0% and 3% of the cases.

**Table 5.** Space efficiency of the encoding schemes of CQF and CQF-deNoise. This table shows the numbers of counter values *C* within the given ranges for which the encoding scheme of CQF-deNoise requires one fewer (−1), the same (0) or one more counter slot, considering all possible remainders that contain *r* = 8 bits.

| Range of $C$ | Number of CQF-deNoise slots - number of CQF slots | | |
| --- | --- | --- | --- |
|  | -1 | 0 | 1 |
| $[1, 2^8]$ | 40,255 (61%) | 25,279 (39%) | 2 (0%) |
| $[1, 2^{16}]$ | 2,731,697 (22%) | 12,541,391 (75%) | 504,128 (3%) |

### The de-noise method: overview

The main idea of our de-noise procedure is to identify low-frequency k-mers and remove them from the CQF during the counting process. The objectives are: 1) to remove false k-mers as early as possible, and 2) to avoid wrongly removing true k-mers. Intuitively, false k-mers can be more confidently identified at the late stage of counting, since at that time the true k-mers should have clearly higher counts than the false ones. In contrast, at the early stage of counting, even true k-mers may also have low counts, making them indistinguishable from the false k-mers. There is thus a trade-off between our two objectives. We handle it by taking a user-specified parameter of the tolerable ratio of true k-mers being wrongly removed, to determine the number of rounds of noise removal and the suitable time for performing each round.

Specifically, CQF-deNoise removes suspected false k-mers *m* times during the counting process, including one round at the end of counting. In each round, all singleton k-mers (i.e., those having an occurrence count of one at that time) are considered the suspected false k-mers. As a result, any k-mer with an occurrence count larger than *m* is guaranteed to remain in the CQF at the end of the counting process, although its final count can be smaller than its original count by a difference up to *m* − 1. On the other hand, k-mers with an occurrence count equal to or smaller than *m* may or may not remain in the CQF at the end of the counting process, depending on whether it appears exactly once between every two rounds of noise removal.

The number of noise removal rounds, *m*, is determined as follows. First, define *m*^′^ as the largest integer such that the fraction of true k-mers with an occurrence count of *m*^′^ or less is smaller than the user specified threshold. In other words, *m*^′^ serves as a conservative estimate of the maximum number of noise removal rounds that can be performed, and it can be estimated based on the genome size, sequencing depth and sequencing error rate. On the other hand, as to be explained below, the size of the CQF depends on the number of noise removal rounds, and different numbers of rounds could lead to the same CQF size. For instance, if *m*^′^− rounds and *m*^′^− 1 rounds would both lead to the same CQF size, it is better to use *m*^′^ 1 rounds because the rate of wrongly removing true k-mers would be smaller but the memory requirement stays the same. Therefore, the actual number of rounds of noise removal, *m*, is chosen as the smallest integer such that the CQF size would be the same as the one with *m*^′^ rounds of noise removal.

### The de-noise method: estimating the true-to-false k-mer ratio

Suppose the genome size is *G* and sequencing reads each of length *l* have been generated to an average genome-wide depth-of-coverage of *d*. The total number of k-mers on these reads is *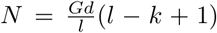*. Suppose that among these k-mers, the ratio of true k-mers that come from the genome to false k-mers that occur due to errors is *R*, the number of true k-mers will be 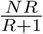. Finally, if the number of unique true k-mers is *u*, their average occurrence count will be 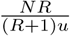. Accordingly, the occurrence counts of the true k-mers are expected to follow a Poisson distribution with parameter *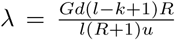* and *m*^′^ = ⌊*F*^*−*1^(*w*)⌋, where *F* is the cumulative distribution function of the Poisson distribution and *w* is the user-specified tolerable ratio of true k-mers being wrongly removed.

In the above formula, *l* and *d* are properties of the sequencing data, *k* and *w* are user parameters, *G* is either prior knowledge supplied by the user or estimated using an efficient method such as ntCard [29], which provides basic statistics of k-mers but cannot give the occurrence counts of individual k-mers. The total number of unique true k-mers, *u*, is not known *a priori*, but an upper bound of it can be used, the value of which can be obtained by assuming all k-mers in the genome are unique. This leaves us with the last variable, the true-to-false k-mer ratio, *R*.

Here we describe an algorithm that can compute this ratio based on the error profile of sequencing reads, i.e., the base error rate of each read position. The error profile is platform dependent. For example, for Illumina short reads, the base error rate is highest at the beginning and at the end of each read. If the error profile is not available, one may instead assume a constant base error rate at each read position, in which case the true-to-false k-mer ratio can be computed using a simple formula as we will see below. For simplicity, we assume that whenever an error occurs in a k-mer, the resulting k-mer always does not exist in the genome, although in reality a small portion of these error-containing k-mers can actually be found in the genome.

The main difficulty of this calculation is that there are spatial dependencies in two ways. First, if a base error appears at a read position, it turns all k-mers that overlap this position into false k-mers at the same time. Second, if multiple base errors appear at nearby positions, together they create a smaller number of false k-mers (with some false k-mers containing multiple errors) than when they are far apart.

To handle these spatial dependencies, we use a dynamic programming algorithm to compute the probabilities of error positions. Suppose the error profile is given in the form of a length-*l* vector of base error probabilities *e*, where *l* is the length of each sequencing read. We define a two-dimensional table *V* (*i, j*) to denote the probability that among the *k* consecutive positions ending at position *i* (where *i* ranges from *k* to *l*), the *j*-th of them is the last position with a base error. For example, if *k* = 4, *V* (6, 2) is the probability that among read positions 3, 4, 5 and 6, the last error occurs at position 4 (which is the second position among these positions). We also define *V* (*i*, 0) as the probability that among the *k* consecutive positions ending at position *i*, there is not a base error.

Table *V* is initialized by considering the first row, *i* = *k*:

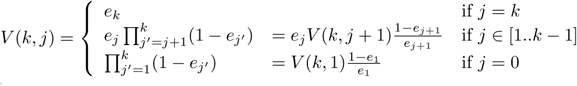

The remaining rows of *V* can be filled in according to the values in the previous row:

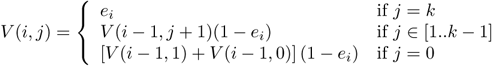

Since we assume that a k-mer is a true k-mer if and only if there is not a base error in any of the k positions, the expected number of true k-mers in a read is 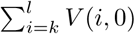 Therefore, the true-to-false k-mer ratio is given by 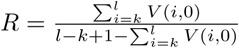

The *V* table contains (*l − k*+1)(*k*+1) entries, each requiring a constant amount of time to fill in. Therefore, the time complexity of the algorithm is *O*((*l* − *k*)*k*), which is also a constant with fixed *k* and *l*. The space complexity is *O*(*k*), since once a row has been filled in, all entries in the previous row can be discarded.

If the detailed error profile is not available and every read position is assumed to have the same base error rate of *e*_0_, it can be easily proved that the true-to-false k-mer ratio is *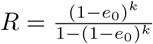*.

To test our algorithm, we generated a simulated data set using ART [30] assuming the Illumina MiSeq v3 protocol. Using the F. vesca reference genome FraVe-sHawaii 1.0, we generated reads of *l*=250bp to an average genome-wide depth-of-coverage of *d* = 50 ×. By aligning the reads to the reference, the average base error rate was found to be 0.4%, and the actual ratio of true k-mers to false k-mers was 8.929 when k was set to 28. When we assumed every position to have the same base error rate of *e*_0_ = 0.4%, we obtained an estimated true-to-false k-mer ratio of *R* = 8.409 based on the above formula. When we instead ran our dynamic programming algorithm using the platform-specific error profile, we obtained an estimated true-to-false k-mer ratio of *R* = 8.873, which is closer to the actual value of 8.929.

If the error profile involves indels, which are common for some platforms such as PacBio SMRT sequencing, our dynamic programming algorithm can be extended to handle additional insertion and deletion states between read positions. We do not further pursue this direction in the current study.

## The de-noise method: time of performing de-noise

With the true-to-false k-mer ratio estimated, the number of rounds of k-mer removal can be computed accordingly as explained. The next question is when the other *m* − 1 rounds should be carried out. Based on the variables defined above, the total number of false k-mers is 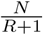 To keep the size of the CQF at its minimum, it would be the best to remove these false k-mers evenly across the *m* rounds of removal, assuming that many of them occur only once in all the reads (and thus they can be removed in any of the *m* rounds). Based on this idea, in the worst case, if all the true k-mers have already been encountered before the target number of false k-mers have been encountered, the CQF will contain *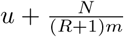* unique k-mers in total. This threshold serves as the trigger to start the next round of k-mer removal.

Although in the above discussion we ignored repeats in the genome, as shown in the Results section, our procedure is effective in removing false k-mers in our empirical tests.

Another limitation of the above derivations is that we used the probability for a true k-mer to have an occurrence count no more than *m* to quantify wrong removals, but such k-mers are not necessarily removed by our procedure, since each of them would not be removed if there are two new occurrences of it between two rounds of removal. Therefore the wrong removal rate was over-estimated.

A more accurate wrong removal rate can be calculated as follows. Suppose there are *m* rounds of removal, which separate all the k-mers into *m* bins with the *i*-th bin containing *b*_*i*_ k-mers. Then in order for a true k-mer with a total occurrence count of *c* ≤*m* to be wrongly removed, these *c* occurrences must be distributed into *c* different bins, and there are 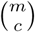 different ways. For the specific way involving bins *i*_1_, *i*_2_, *…, i*_*c*_ where 1 *≤ i*_1_ *< i*_2_ *< … < i*_*c*_ *≤ m*, the probability of happening is assuming the occurrence order of k-mers is random. The wrong removal rate for k-mers having c occurrence counts can then be computed by summing 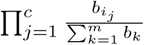 the probabilities. This rate for k-mers having *c* occurrence counts can then calculation is feasible only when the number of k-mer removal rounds is small, and is thus not implemented in CQF-deNoise.

### Comparing CQF-deNoise with other k-mer counting methods

We compared the running time and memory usage of CQF-deNoise with four other k-mer counting methods using four data sets with diverse properties (Table 1). All our tests were run on a machine with Intel(R) Xeon(R) CPUs (E7-4850 v3 @2.20GHz with 112 cores and 35.8MB L3 cache) and 504GB RAM. All programs were run with 16 threads. Running time was defined as the wall clock time, during which the program loaded and parsed the sequencing data, counted the k-mers, and wrote the results to output files on the disk. Memory consumption was defined as the peak resident set size (RSS).

All methods were tested for *k* = 28 and *k* = 55, which are values also used in some previous studies [13, 15]. To handle the issue that sequencing reads could come from either strand, among each k-mer and its reverse complement, we converted the one with a larger hash value to the one with a smaller hash value before counting.

To make the comparisons fair, we ran all programs with the option of removing singleton k-mers chosen if this option was provided as follows. For BF-Counter, it had the ability to remove singleton k-mers by only counting the non-singletons in the second structure. For CQF-deNoise, we set the wrong removal rate tolerance threshold *w* to the conservative value of 1/genome-size, and let the algorithm determine the number of rounds of noise removal automatically. We found that the number of noise removal rounds remained unchanged for long ranges of this threshold value (Table 6), showing that the counting results would be highly stable for different values used. For Jellyfish2 and KMC3, we selected the mode to save k-mers with an occurrence count at least 2. Squeakr did not provide an option for removing singleton k-mers. Since a brute-force enumeration of all the k-mers and removal of the singletons after counting could be very slow, we did not perform it.

**Table 6.** Statistics for running CQF-deNoise on 4 dataset with k=28. The wrong removal rate tolerance thresholds were automatically determined by CQF-deNoise. The ranges of wrong removal rate tolerance thresholds show the values of this parameter used that would lead to the same number of noise removal rounds.

| Species | q | Estimated mean coverage of true k-mers | Wrong removal rate tolerance threshold | Number of noise removal rounds | Range of wrong removal rate tolerance thresholds leading to the same number of noise removal rounds | Actual wrong removal rate of true k-mers |
| --- | --- | --- | --- | --- | --- | --- |
| C. elegans | 29 | 171 | $8.4 \times 10^{-9}$ | 17 | $[1.6 \times 10^{-51}, 1]$ | $1.6 \times 10^{-51}$ |
| F. vesca | 31 | 29 | $2.9 \times 10^{-9}$ | 2 | $[1.1 \times 10^{-10}, 7.3 \times 10^{-4}]$ | $1.1 \times 10^{-10}$ |
| Z. mays | 33 | 60 | $6.2 \times 10^{-10}$ | 1 | $[5.3 \times 10^{-25}, 6.9 \times 10^{-6}]$ | $5.3 \times 10^{-25}$ |
| H. sapiens | 33 | 61 | $3.7 \times 10^{-10}$ | 11 | $[4.3 \times 10^{-15}, 1]$ | $4.3 \times 10^{-15}$ |

BFCounter and Jellyfish2 required the maximum number of unique k-mers as input, which we computed using ntCard [29]. CQF-deNoise required the number of true k-mers as an estimation of genome size and the total number of k-mers as inputs, and Squeakr required the number of slots in the CQF as input, which were all estimated based on the output of ntCard. CQF-deNoise also used the results of ntCard to determine the fraction of singleton k-mers for estimating the base error rate *e*_0_ in the comparisons. In the time measurements, the running time of ntCard was also added to the total running time of Squeakr and CQF-deNoise but not BFCounter or Jellyfish2, since the latter two methods could also obtain their required inputs by some faster means.

Squeakr could use x86 bit manipulation instructions to speed up its counting by a factor of 2-4 [13]. Our machine did not support these instructions. Although the resulting running time could not fully reflect the counting speed of Squeakr, CQF-deNoise was also disadvantaged in the same way, because our implementation could also use these instructions. When running Squeakr, we used the fast ntHash function [28] rather than the default Murmur hash function.

For all other parameters of the k-mer counting methods, we used their default values.

### K-mer counts-based clustering of single cells

For 10x scRNA-seq data, we considered two strategies for selecting the high-quality cells to analyze. In the first strategy, the *N* cells (identified by their barcodes) with the highest UMI counts are selected, where *N* is a user parameter. The second strategy, adopted from CellRanger, first computes the 99th percentile of the UMI counts in the top *N* cells. It then selects all cells with an UMI count at least one-tenth of that value for the analysis. Both strategies are provided as user options in our implementation. On the other, for the particular 10x data set we analyzed in this study, we considered only the 713 cells selected by CellRanger for a fair comparison of the two methods.

We compared two approaches to defining k-mer count profiles of single cells. In the first approach, all k-mers on the scRNA-seq reads were counted. In the second approach, only the k-mers on protein-coding transcripts were counted, and the “genome size” was defined as their total length. To use the second approach, we first used the gffread utility in cufflinks (v2.2.1) [31] with options “-CME” to extract 120,463 mRNA sequences with CDS features from the Ref-Seq human gene annotation corresponding to reference genome GRCh38.p12. We then extracted all the k-mers on these sequences and stored them in a Bloom filter. For each k-mer on the scRNA-seq reads, we first checked its presence in the Bloom filter, and inserted into the CQF only if it was found.

For both approaches, after k-mer counting we first normalized each counter by dividing it by the total count in the whole CQF. We then computed the distance between every two cells as the sum of the absolute difference of these normalized counts of every object in the two CQFs, where the count of a non-existing object was defined as 0 (when the object was only contained in the CQF of the other cell but not this one). Finally, for each cell we kept only the distance values of its five nearest neighbors and set all the other distance values to the maximum value of one, such that the final distance measure corresponds to one that comes from a weighted nearest-neighbor graph.

In our implementation, both k-mer counting and the distance matrix calculation can be run with multiple threads in parallel.

To benchmark the k-mer-based analysis results, we ran CellRanger (v3.0.2) to produce the two-dimensional projection of cells and the cell clusters. The options “–localcores” and “–localmem” were used. We used SingleR (v1.0) to annotate the cell types of single cells with default settings. For our k-mer-based approach, it took the distance matrix as input. For CellRanger, it took the filtered feature matrix as input.

### Additional tools provided in our implementation

We provide a list of additional tools for various CQF operations, including downsizing, intersection, addition, and subtraction, which are useful in different applications. These tools enable logical and arithmetic operations to be directly performed on the object occurrence counts efficiently rather than the raw sequencing reads that would require a lot more time.

Downsizing is to resize a CQF that uses *p*_1_ bits of hash value to one that uses *p*_2_ (*p*_1_ *> p*_2_) bits. One use of it is to compress the data structure to use less space, especially when the occupancy is low. It also facilitates other operations such as taking the intersection or difference of two CQFs. Downsizing can be efficiently performed by simply iterating through all the objects in the original CQF and inserting the transformed hash values into the new CQF, keeping only the first *p*_2_ bits of its original signature as its new signature. Since objects originally having the same quotient will still have the same quotient after the downsizing, their sort order is maintained, which avoids shifting of the contents, the part usually most time consuming during object insertion.

Intersection is to identify the objects commonly contained in two CQFs and produce two new CQFs that contain only the common objects and their corresponding counts in the original CQFs.

Addition and subtraction respectively compute the sum and difference of the object occurrence counts in two CQFs. In the case of subtraction, objects with a resulting negative count are removed from the output CQF.

## Code availability

Our implementation of CQF-deNoise is available at https://github.com/Christina-hshi/CQF-deNoise.git under the BSD 3-Clause license.

### Author’s contributions

CHS and KYY conceived the study and designed the methods. CHS implemented the methods and performed the empirical studies. CHS and KYY analyzed the data and prepared the manuscript.

## Acknowledgements

We thank Jianquan Cao and other members in the Yip Lab for helpful discussions. KYY was partially supported by the Hong Kong Research Grants Council General Research Fund 14170217, Collaborative Research Funds C4054-16G, C4045-18WF, C4057-18EF and Theme-based Research Scheme T12C-714/14-R. This work was also supported by the Hong Kong Epigenomics Project (EpiHK).

